# Pathogen-prey-predator relations of avian raptors during epizootics of highly pathogenic avian influenza virus HPAIV H5N1 (clade 2.3.4.4b) in Germany

**DOI:** 10.1101/2023.11.19.567176

**Authors:** Anne Günther, Oliver Krone, Anja Globig, Anne Pohlmann, Jacqueline King, Christine Fast, Christian Grund, Christin Hennig, Christof Herrmann, Simon Piro, Dennis Rubbenstroth, Jana Schulz, Christoph Staubach, Lina Stacker, Lorenz Ulrich, Ute Ziegler, Timm Harder, Martin Beer

## Abstract

Transition of highly pathogenic clade 2.3.4.4b H5 avian influenza virus (HPAIV) from epizootic to enzootic status in Northern European countries was associated with severe losses and even mass mortalities among various wild bird species. Both avian and mammalian raptors hunting infected debilitated birds or scavenging on virus-contaminated avian carcasses contracted HPAIV infection. This precarious pathogen-prey-predator relation further worsened when in 2021 and 2022 outbreaks in Germany overlapped with the hatching season of avian raptor species. Retro- and prospective surveillance revealed avian raptors as important indicators of HPAIV and its genetic diversity on the one hand. On the other hand, their role as victims of HPAIV is stipulated. The first case of an HPAIV H5N1-related death of a white-tailed sea eagle (*Haliaeetus albicilla;* WTSE) hatch in Germany, 2021, followed by several such cases in 2022, and a low overall seropositivity rate of 5.0-7.9% among WTSE nestlings, raised fears of a serious negative impact on reproduction rates of WTSEs and other birds of prey when HPAIV becomes enzootic in an ecosystem. However, comparably stable breeding success of WTSE in the study area in 2022 and a potentially evolving natural immunity raises hope for a less severe long-term impact.

**Article impact statement:** Adapted surveillance measures were developed to assess risks for the conservation of avian raptors due to the panzootic spread of HPAIV.

## Introduction

The current panzootic of highly pathogenic (HP) goose/Guangdong (gs/GD) clade 2.3.4.4b avian influenza virus (AIV) causes immense damage in poultry holdings and severe die-offs in wild birds worldwide (Caliendo, Lewis, et al., 2022). Along with an enormous extension in geographic range, the recent gs/GD HPAIV H5 lineage has gained an enzootic status in European wild bird populations (A. Pohlmann et al., 2022) causing immense clinical impact and high mortality in several endangered wild bird species.

The natural history of influenza A viruses (IAV) of low pathogenicity (LPAI) identifies wild water birds of the Anseriformes and Charadriiformes as reservoir hosts. Extended co-evolution ensured efficient virus replication and spread while not impacting the clinical status of the avian hosts (Globig et al., 2013; Globig et al., 2009; Yoon, Webby, & Webster, 2014). However, species of these orders are equally susceptible, and clinically highly vulnerable, to HPAIV which arise sporadically by spontaneous mutation in galliform poultry infected with LPAI precursor viruses of subtypes H5 or H7 (Pantin-Jackwood & Swayne, 2009). During the 1990s, such HPAIV (i.e. the gs/GD lineage) arose in Chinese poultry populations and reached migratory wild bird populations by spill-over infections in Far East Asia since the early 2000 years. In fact, migratory waterfowl has been identified as long-distance vectors of gs/GD HPAIV H5 (Global Consortium for H5N8 and Related Influenza Viruses (2016), 2016). Along with HPAIV dissemination in wild water birds, avian raptors of the orders Accipitriformes, Falconiformes and Strigiformes are increasingly affected (EFSA (European Food Safety Authority), ECDC (European Centre for Disease Prevention and Control), EURL (European Reference Laboratory for Avian Influenza), Adlhoch, Fusaro, Gonzales, Kuiken, Marangon, Niqueux, Staubach, Terregino, Aznar, Chuzhakina, et al., 2022; EFSA (European Food Safety Authority), ECDC (European Centre for Disease Prevention and Control), et al., 2022a, 2022b; EFSA (European Food Safety Authority), ECDC (European Centre for Disease Prevention and Control), EURL (European Reference Laboratory for Avian Influenza), Adlhoch, Fusaro, Gonzales, Kuiken, Marangon, et al., 2023). Due to the high public attention that many of these species receive, their feeding behaviour on diseased and weakened prey or infected carcasses and their apparently high susceptibility, they were marked out as indicator species for (passive) HPAIV disease surveillance (Caliendo, Leijten, van de Bildt, Fouchier, Rijks, & Kuiken, 2022; El Zowalaty et al., 2022; Günther et al., 2022; Krone et al., 2018; Nemeth et al., 2023; Redig & Goyal, 2012; van den Brand et al., 2015).

The recently established year-round presence of gs/GD HPAIV H5 in European wild bird populations poses major threats to avian raptors: (i) Increased infection pressure due to multiple opportunities of ingesting HPAIV H5 infected prey (Banyard et al., 2022; Anne Pohlmann et al., 2023; Rijks et al., 2022) and (ii) a temporal overlap of virus presence with the hatching season of raptor chicks. Mortality among nestlings of white-tailed sea eagles (*Haliaeetus albicilla*, WTSE) in Estonia in 2021 (Estonian University of Life Sciences, 19.05.2021) and bald eagles (*Haliaeetus leucocephalus*) in North America in 2022 due to alimentary HPAIV H5 infections (Nemeth et al., 2023) have been reported already.

Within a nationwide retrospective and regional prospective surveillance for HPAIV H5 infections in raptor species in Germany we: *(i)* report on (HP)AIV infection rates in raptors since 2016, (ii) screened archived samples of avian predators collected across Germany since 2010, and (iii) prospectively sampled raptor nestlings during ringing activities in Mecklenburg-Western Pomerania (MWP), Germany. This region, severely affected by gs/GD HPAI in 2021-22, holds the highest density of WTSE breeding pairs in Germany and harbours important stop-over sites for migratory water birds (Herrmann, Krone, Stjernberg, & Helander, 2023; Krone et al., 2018).

## Material and Methods

### Sample and data sets

All samples obtained in a prospective or retrospective surveillance approach were collected in Germany including individuals from the taxonomic orders of Accipitriformes, Strigiformes and Falconiformes. For reasons of endangered species protection listing precise breeding locations was omitted throughout this manuscript.

### Retrospective surveillance on HPAIV in raptor species

The avian influenza database represents a governmental, non-public database on all virological data regarding AIV infections in wild birds. Data were selected with respect to the orders Accipitriformes, Falconiformes and Strigiformes, subsequently considered as raptors, and on HPAI-specific results on March 6, 2023, for the years 2016 to 2022, covering the activity of HPAIV clade 2.3.4.4b strains.

The database survey was compiled by raptor samples archived by the Leibnitz Institute for Zoo and Wildlife Research, Berlin. These organ samples (mainly brain, lung or liver) had not been examined for HPAIV previously as they were collected in frame of unrelated research projects. The carcasses were collected across the whole geographic range of Germany. All samples were examined at the Friedrich-Loeffler-Institut, Isle of Riems, Germany. Additionally, WTSE sera retrieved as part of different research projects in the federal states of Brandenburg, MWP and Thuringia were included.

### Prospective surveillance in raptor nestlings and rehabilitated raptor species

The majority of individuals was sampled as nestlings of ten different species, when handled within their first weeks of age during scientific bird ringing activities in spring 2021 (April to July) and 2022 (May and June) in MWP, Germany (see 2.1.3 for permissions). The ringing of birds allows for an unambiguous and unmistakable individual identification of an animal (and thus a sample) from that point on. Some birds were sampled in a wild bird rescue centre in Greifswald, MWP (June, July and October 2021 and May and June 2022), covering five different species. The sample-identification comprises serial numbers indicating the affiliation to a nest/location, while letters represent the sampled individuals per sampling nest/location (e.g., two individuals at location (nest) #6 are named #6A and #6B). All birds were physically examined for general behaviour and clinical signs of infection, e.g. laboured breathing or neurological disease manifestation. A complete sample set included two separate swabs (oropharyngeal and cloacal) and a venous blood sample, taken from the wing vein. In some cases, only a subset of samples was taken, either to reduce the time of handling, due to situation-dependent field-work aspects, or according to the bird’s size or clinical condition.

### Ethical statement

Organ samples from raptor carcases were collected during post-mortem examinations in the context of different research projects with ecotoxicological objectives and, therefore, no additional permits were required for our retrospective analyses. The serum samples collected in prior studies were approved by the authority of the Federal State of MWP, Germany (LALLF reference number 7221.3-3.2-004/19) and by the authority of the Federal State of Brandenburg (LAVG reference number 2347-A-10-1-2019).

The prospective sampling of avian raptors in MWP, Germany, was approved by the authority of the Federal State of MWP, Germany (LALLF reference number 7221.3-2-003/21, approved 24 March 2021).

### Molecular analyses

Swabs were stored in virus cultivation medium (Sigma-Virocult®). Archived swabs and organ samples were kept at-70 °C until final analyses. RNA extraction from swabs and supernatants of homogenated organ samples was performed using the Macherey-Nagel NucleoMag® VET-Kit on a KingFisher Flex Purification System (Thermo Fisher Scientific), following the manufactures’ instructions. A heterologous internal control RNA was added during the RNA extraction process (B. Hoffmann, Depner, Schirrmeier, & Beer, 2006), to assure successful extraction process. RNA was screened by real-time reverse transcription polymerase chain reaction (RT-qPCR) for presence of IAV-specific generic targets in matrix (M) or nucleoprotein (NP) genes (Fereidouni et al., 2012; E. Hoffmann, Stech, Guan, Webster, & Perez, 2001). Positive samples were further sub- and pathotyped by RT-qPCR protocols as described previously (Hassan et al., 2022).

### Serological analyses

Samples of coagulated blood were transported cooled and dark until separation from serum and blood cruor by ten minutes of centrifugation (3500 rpm). Serum was stored at-20 °C after heat inactivation for 30 minutes at 56°C. Sera were screened using competitive enzyme-linked immunosorbent assays for IAV-specific antibodies. In a first step, all samples were applied to the ID Screen® Influenza A Antibody Competition Multi-species assay, detecting generic antibodies against the NP. In case of positive findings, those samples were screened by using the ID Screen® Influenza H5 Antibody Competition assay to detect antibodies against the HA of subtype H5. The cut-off values for sample to negative (S/N) ratios were used as recommended by the manufacturer: S/N%≤45% positive, 45<S/N%<50 indeterminate and S/N%≥50 negative for antibodies against NP, respectively S/N%≤50% positive, 50<S/N%<60 indeterminate, S/N%≥60 for H5. Due to limited sample volumes, a single test per sample and step was performed. A single sample, for which sufficient volume was available, was additionally analysed in a hemagglutinin inhibition (HI) test against a set of reference antigens supplied by the European Reference Laboratory Padova, Italy (H5N1, Eurasian AIV: A/ck/Scotland/1/59; H5N3, Eurasian AIV: A/Teal/England/7394-2805/06; H5N8, HPAIV gs/GD: A/tk/Italy/7898/14; Newcastle disease virus Clone 30).

### Sequencing and genetic analyses

HPAIV-positive samples were considered for sequencing when revealing distinct viral loads of Cq (quantification cycle)-values below 30. The sequencing workflow described by King, Harder, Beer, and Pohlmann (2020) was followed. Retrospective sequences from other studies and from databases were included for comparison and genotype assignment. Genotype differentiation and derivation of reference sequence were done with a combined phylogenetic and similarity-based method. Genotypes were assigned, and new genotypes were differentiated if they are clustering separately with robust bootstrapping values (>80) or if differences greater than 2% were observed when comparing nucleotides at segment level. The first complete genome sequence of a newly detected genotype was used as a reference sequence, and additional references for a genotype were derived as needed. Genotype names are filed to include locality (three digits), date of first discovery (month-year), and NA subtype. When multiple genotypes of one subtype were assigned within the same locality and date, the names were numbered consecutively. Detailed methodology and overview of reference sequences are available as technical note under https://doi.org/10.5281/zenodo.8233814.

### Breeding success rate and breeding pair numbers of white-tailed eagles in MWP, Germany

We utilized data on the breeding success rate and the number of overall breeding pairs for WTSEs in MWP, from 2002 to 2022. These data sets were compiled by the “Working Group for Conservation of Large Birds MWP” and provided by the Agency for Environment, Nature Conservation, and Geology MWP.

### Statistical analyses

For the calculation of the 95% confidence intervals (95%CI) (Clopper & Pearson, 1934) and the Fishertest (Fisher, 1936) we applied R version R4.2.2 (R Core Team, 2021). The 95%CI is provided for the detection rate of (HP)AIV RNA positive species or groups of species. We utilized the Fisher-test to verify, if there is a significant difference between the findings on NP-specific antibodies in WTSE nestlings and all other sampled raptor nestlings (significant value is considered as p<0.05).

## Results

### Retrospective sample screening confirms large to medium-sized raptors highly affected by HPAIV H5

In a nationwide retrospective surveillance organ samples from 232 birds of ten different species collected between 2010-2022 were analysed retrospectively: Ospreys (n=2; *Pandion haliaetus*), Northern goshawks (n=1; *Accipiter gentilis*), common buzzards (n=46; *Buteo buteo*), red kites (n=16; *Milvus milvus*), barn owls (n=23; *Tyto alba*), common kestrels (n=28; *Falco tinnunculus*), tawny owls (n=28; *Strix aluco*) and peregrine falcons (n=3; *Falco peregrinus*), WTSE (n=82) and Eurasian eagle owls (n=3; *Bubo bubo*). Different age cohorts, from nestlings to adult individuals, were represented (Figure 1).

**Figure 1.**
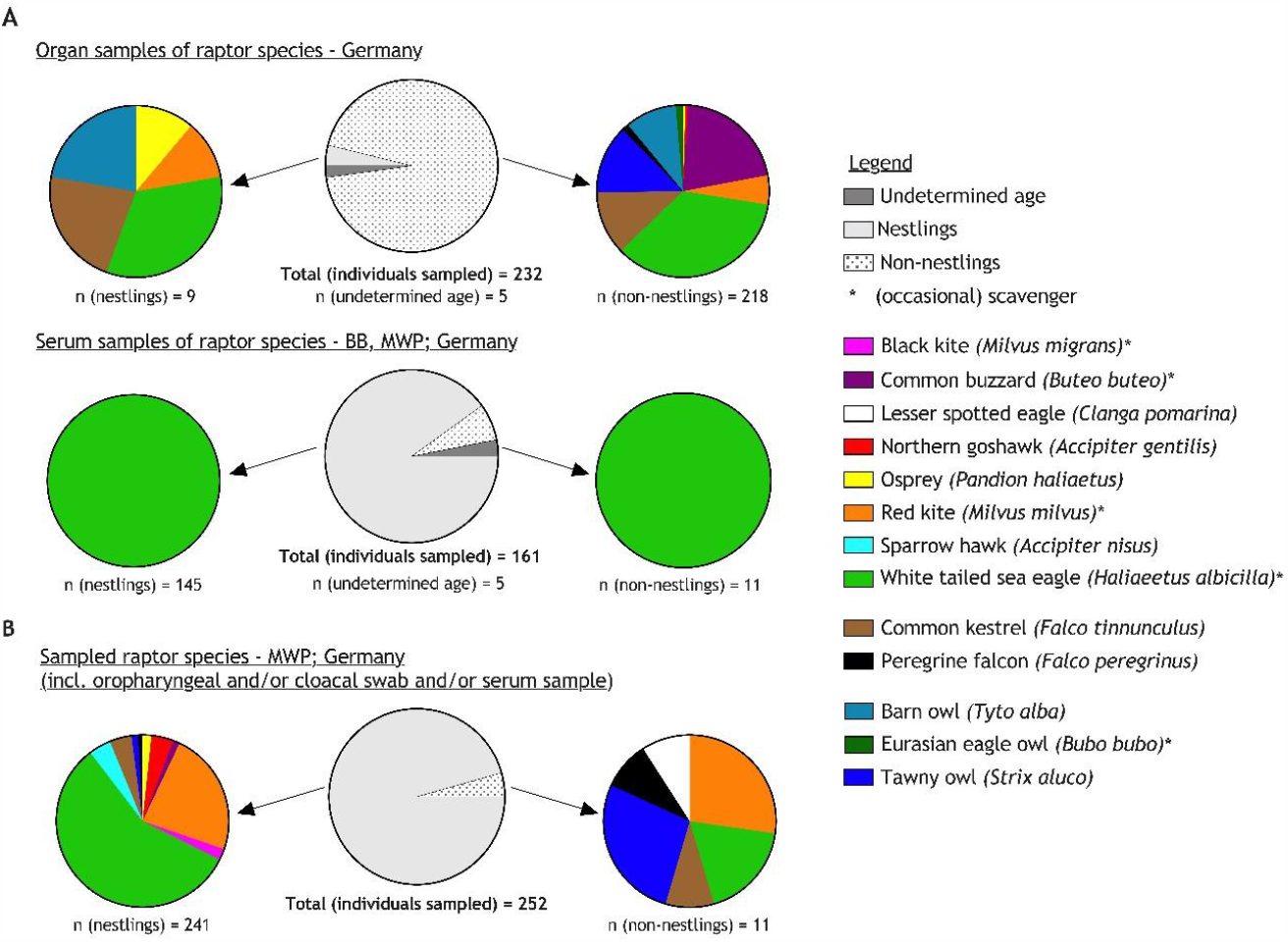
Overview of individual avian raptors sampled within the retrospective (A) or prospective (B) surveillance approach in this study, including additional information on age cohorts and scavenging behavior.

The general German wild bird surveillance revealed yearly HPAIV H5 detection rates between 0.0% (95%CI 0.0-2.3) and 7.6% (95%CI 5.2-10.6) in raptors for the years 2016 to 2022 (Figure 2). The highest detection rate of HPAIV-positive raptors is found in WTSEs (13.3 %; 95%CI 8.79-19.00), buzzard sp./common buzzards (6.55%; 95%CI 4.43-9.27 and 4.75%; 95%CI 3.69-5.99, respectively), Northern goshawks (4.6%; 95%CI 2.23-8.31) and peregrine falcons (3.89%; 95%CI 1.27-8.81) – followed by other raptor species mentioned in the supplementary material Table S1. Two of the HPAIV H5-positive WTSE samples from 2021 were identified as nestlings from a single breeding location in Schleswig-Holstein (SH), Germany.

**Figure 2.**
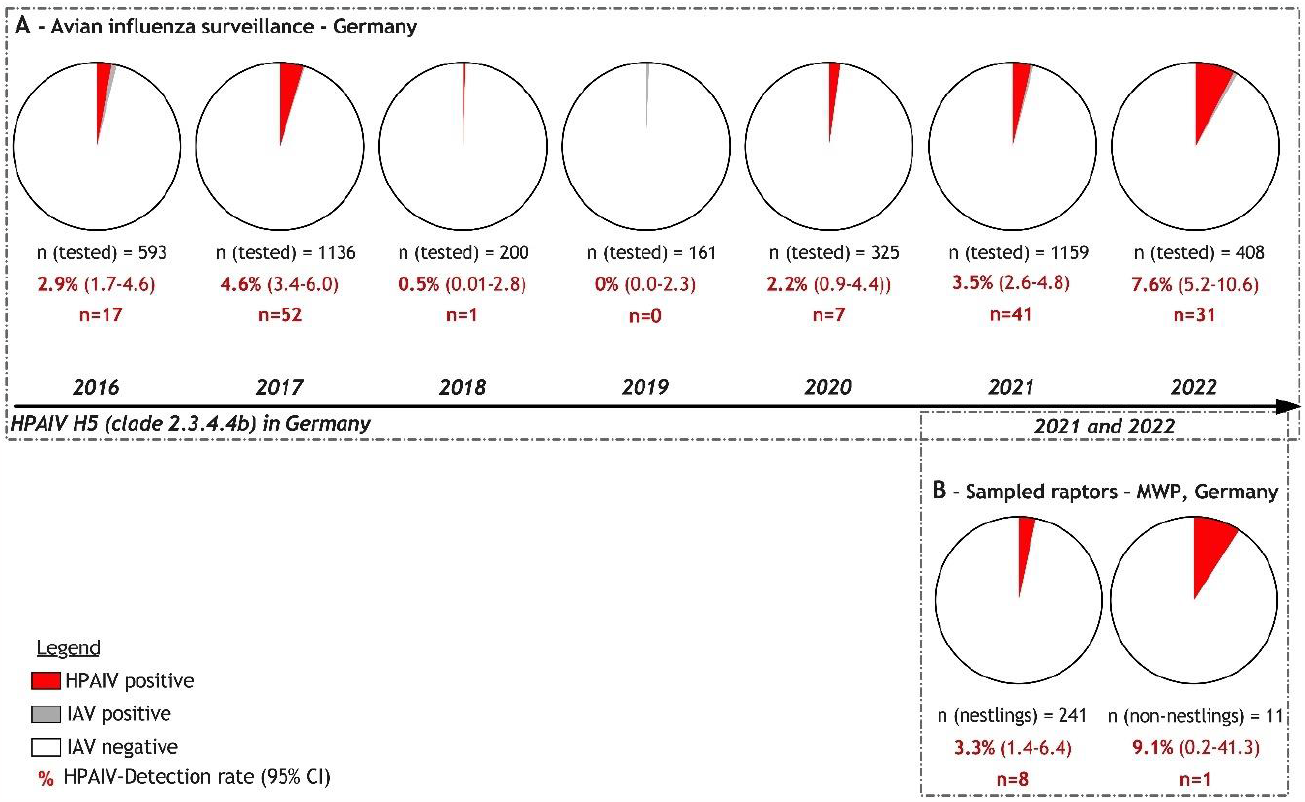
Numbers of raptors screened for influenza A viruses (IAV) and testing positive for gs/GD highly pathogenic avian influenza viruses (HPAIV). The proportion of HPAIV-positive birds is given for the retrospective surveillance across the whole of Germany between 2016-2022 (A) and nestlings and non-nestlings (B), sampled within the prospective-regional sampling approach in Mecklenburg-Western Pomerania (MWP), a Federal State in the Northeast of Germany (2021 and 2022).

### Prospective surveillance in WTSE and their nestlings revealed increased HPAIV H5N1 infection rate since 2021

A prospective surveillance of nestlings started in early 2021 and was carried out in the context of scientific bird ringing. 252 individual birds of eleven different species were sampled (Figure 1). The majority of samples was obtained between April to July 2021 (n=118) and May to June 2022 (n=124) from nestlings on their nests in natural habitats in the German Federal State of Mecklenburg-Western Pomerania (MWP), Germany. Additionally, ten birds (2021: n=7, 2022: n=3) were sampled in a wild bird rescue centre, of which seven birds were considered as adults and three as fledglings/juveniles. The majority of the nestlings showed no clinical signs. However, for few birds (n=13) healed injuries (red kite, n=1 and lesser-spotted eagle [*Clanga pomarina*], n=1; both adult), poor nutritional status (red kite, n=1, nestling and WTSE, n=1, adult) and increased agitation associated with capture/handling (WTSE, n=2, nestlings) were noted. Two WTSE nestlings appeared mildly (n=1) or markedly depressed (#84A; n=1), and five nestlings showed mild serous rhinorrhoea. During fieldwork, nine WTSE nestlings were found dead, either on the nest or in close proximity under the eyrie in varying states of decay. One of them had been sampled alive approximately two weeks prior to death (#84A). In October 2021, a juvenile WTSE (#72A) was found with neurological disorders (e.g. ataxia) and unable to fly. It was sampled by a veterinarian, including a blood sample for confirmative diagnosis of an expected lead intoxication. The bird died on the following day. A detailed overview on all samples is provided in Tables S2-S4.

As compared to the nationwide surveillance, detection rates for the prospective-regional sampling approach in MWP, Germany, in 2021 to 2022 for HPAIV H5-positive individuals ranged from 3.3% (95%CI 1.4-6.4; n=241) in nestlings to 9.1% (95%CI 0.2-41.3; n=11) in non-nestlings. This is based on the examination of a total of 252 oropharyngeal and 230 cloacal swab samples from 252 individual birds. All samples collected from ospreys (n=4), Northern goshawks (n=10), common buzzards (n=3), red kites (n=59), the lesser-spotted eagle (n=1), black kites (n=5; *Milvus migrans*), sparrow hawks (n=10; *Accipiter nisus*), common kestrels (n=11), tawny owls (n=6) and peregrine falcons (n=3) remained negative. In contrast, nine out of 140 WTSEs were confirmed positive for HPAIV H5N1 (clade 2.3.4.4b). Of these, eight samples were obtained from nestlings, sampled in spring 2022 (Figure 2). Two of these were sampled when nestlings were alive (#77A and 84A), whereas nestlings #77B, 79A, 79B, 84B, 140A and 141A were found dead (Figure 3). HPAIV H5N1 was detected not only in swabs but also in organ samples of these six carcasses. Animal #84A was found dead two weeks after ringing, but its carcass was excluded from the necropsies, due to advanced decay. In addition, a juvenile WTSE (#72A) showing neurological disorders before death tested positive in swabs and organ samples (Figure 3). Highest viral genome loads were found in brain, heart, lung and liver samples (Figure 3).

**Figure 3.**
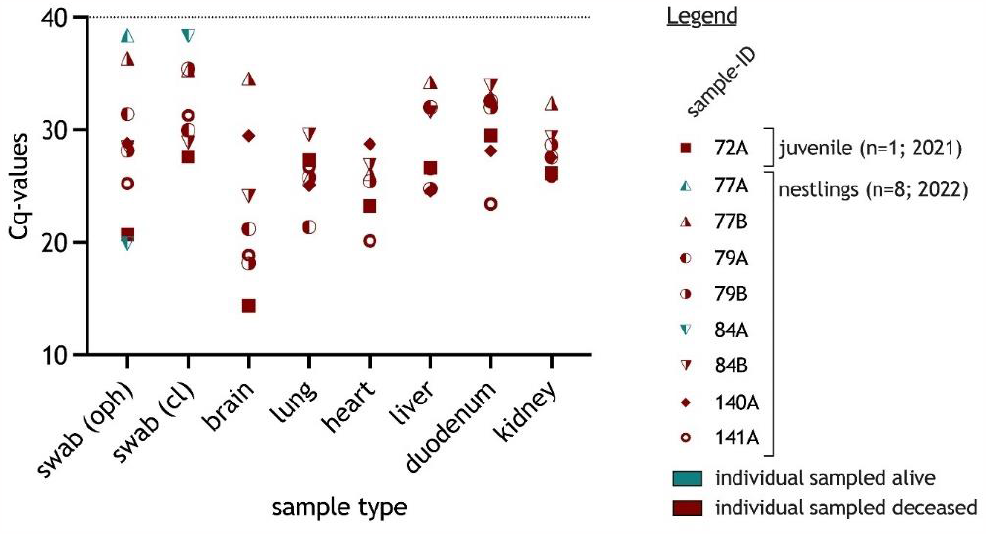
Distribution of relative viral load (low Cq values indicate high viral loads) in HPAIV H5N1-positive white-tailed sea eagles, screened by RT-qPCR. The results are presented per sample matrix (oropharyngeal or cloacal swab or organ) and individual bird. Individuals are assigned numbers and letters, where numbers indicate a specific nest and letters the different nestling therein. Samples taken from individuals alive are shown in blue, samples taken from dead nestlings are shown in red.

### Passive surveillance in avian raptor species in Germany partially mirrors regional diversity of gs/GD HPAIV H5 genotypes

Further virological characterization work and genome-wide sequence analyses on samples in the period between calendar week (CW) 44 in 2020 to CW 48 in 2022 comprised a total of 33 HPAIV H5 genotypes in wild and captive birds, as well as in poultry (Anne Pohlmann, 2023). Eight genotypes (Ger-04-21-N1, Ger-10-20-N5, Ger-10-20-N8, Ger-10-21-N1.2, Ger-10-21-N1.5, Ger-12-21-N1.3 and Ger-12-21-N1.4) were also found in raptors (Figure 2). Genotype Ger-11-21-N1.4 was detected in a buzzard and remained the only finding of that genotype in Germany (highlighted in grey, Figure 4).

**Figure 4.**
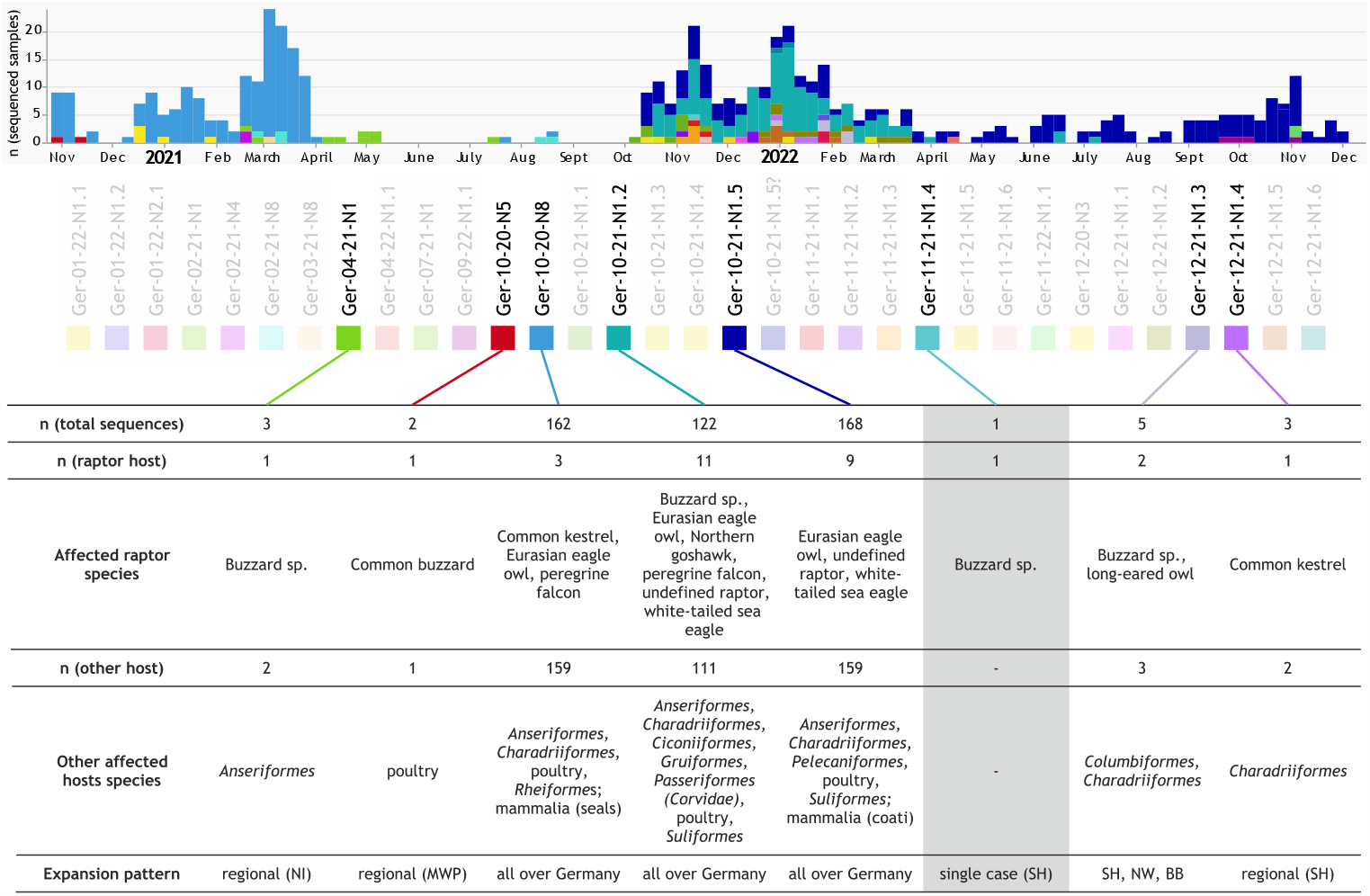
HPAIV H5 genotype variation in Germany. Between calendar week 44 in 2020 and 48 in 2022, 33 distinct genotypes were found in total (unique colour codes, upper panel). Of these, eight were found in avian raptors(highlighted), and additional information on their temporal occurrence and distribution, raptor host species (groups) and other affected hosts is summarized. Ger-11-21-N1.4 is shaded in grey emphasizing that it is the only genotype found exclusively in a raptor host. Abbreviations of the German federal states: BB - Brandenburg; MWP – Mecklenburg-Western Pomerania; NI - Lower Saxony; NW – North Rhine-Westphalia; SH - Schleswig-Holstein.

### Serological evidence of increased AIV, but not H5-specific, exposure rates in WTSE nestlings

In total, 71 (2021) and 114 (2022) serum samples from nestlings of seven different raptor species and seven (2021) and three (2022) serum samples taken from non-nestlings of five different raptor species were prospectively screened for AIV-reactive antibodies (Figure 5-A). Nestlings positive for nucleoprotein (NP)-reactive antibodies were found exclusively for WTSEs (8 out of 116; 6.9%) of which in only one case antibodies against the H5 subtype could be confirmed unambiguously (#103A). In hemagglutinin inhibition (HI) testing this serum revealed the highest titre against a gs/GD-lineage among several H5 antigens of different origins and therefore is highly likely to be clade 2.3.4.4-specific (Table S5). In an adult red kite and WTSE, and in a juvenile WTSE (#72A), NP-antibodies were also detected. H5-specific antibodies could be confirmed for both adult birds, whereas for the juvenile bird (#72A) the result remained indeterminate (Figure 5-A).

**Figure 5.**
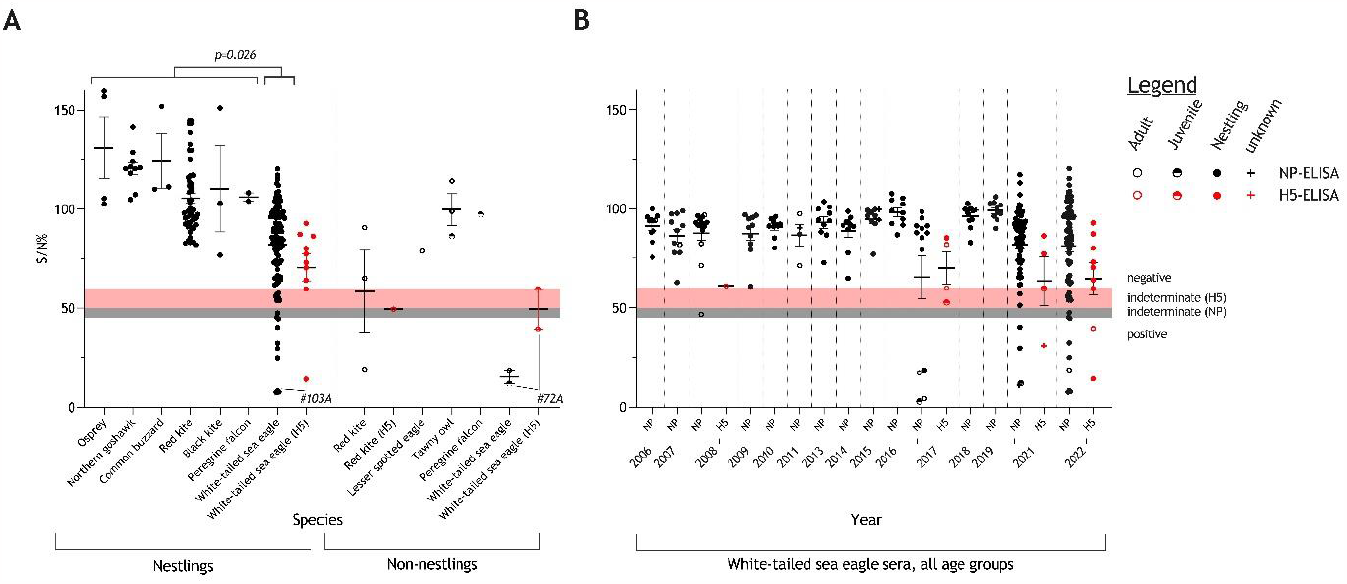
Serological results of samples retrieved from avian raptors. The results are shown in percent inhibition values measured by competition ELISA to detect antibodies against the nucleoprotein (NP, black symbols, grey indeterminate range 45-50%; positive <45%). NP-positive sera were further tested against the hemagglutinin H5 (H5, red symbols, pinkish indeterminate range 50-60%; positive <50%). A) Serological status of individuals sampled within the prospective-regional surveillance approach in Mecklenburg-Western Pomerania (MWP), Germany in 2021 and 2022. The data are grouped per species and age, indicating mean and standard error of the mean. Data points for certain individuals are highlighted by specific numbers; numbers correspond to listings in Supplementary Tables 1-3. P-value < 0.05 is confirming significant differences comparing NP-positive findings in white-tailed sea eagle (WTSE) nestlings (n(positive)=8; n(non-positive)=108) and nestlings of all other raptor species (n(positive)=0; n(non-positive)=69) via the Fisher-test. B) Serological status of WTSE analyzed retrospectively (2001-2019) or obtained within a prospective targeted surveillance approach in MWP, Germany (2021-2). The data are stratified by year of sample origin and by age cohort indicating mean and standard error of the mean.

Furthermore, 161 serum samples from WTSEs, taken during prior investigations in 2006-2011, 2013-2019 and 2021 were retrospectively examined. Those WTSE sera retrieved within prior studies are juxtaposed with serological findings in WTSEs from our prospective surveillance (Figure 5-A). Significantly fewer birds tested seropositive for NP-specific antibodies before 2021, and evidence for H5-specific antibodies was confirmed only in 2021 and 2022 (Figure 5-B).

### No evidence for declining breeding success rate of WTSE in MWP, Germany, despite concurrent enzootic HPAIV H5N1 circulation

As shown in the regions screened in Germany, the amount of WTSE breeding pairs in 2022 was situated in the upper range compared to previous counts over the last two decades (Figure 6-A). The breeding success rate in 2022 when HPAIV H5N1 was highly prevalent in two regions (Isle of Rügen and Isle of Usedom) averaged that of the preceding years (Figure 6-B). The breeding success rate indicates the proportion of those breeding pairs of which at least a single nestling fledged compared to all pairs that had started breeding in the respective year.

**Figure 6.**
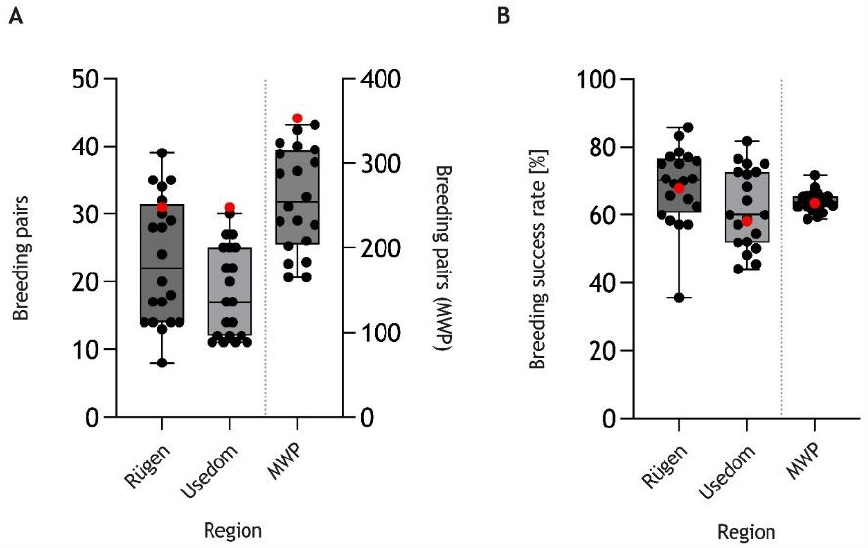
Number of breeding pairs (A) and breeding success rate (B) of white-tailed sea eagles in two selected regions of the federal German state of Mecklenburg-Western Pomerania (MWP; Isle of Rügen and Isle of Usedom) and in total MWP, 2002-2022. Red dots indicate the data for 2022 when HPAIV H5N1 of clade 2.3.4.4b was enzootically prevalent in those regions.

## Discussion

Raptor populations are frequently threatened by a number of anthropogenic factors. These include encroachment of their habitats (Newton, 1979), increased toxicological burdens (Badry et al., 2020; Nadjafzadeh, Hofer, & Krone, 2013) and collision with man-made structures, such as wind energy plants (Heuck et al., 2019), among others. In many countries, including Germany, a high level of conservation effort is required to compensate for these negative anthropogenic factors and has succeeded to stabilize, or even promote growth of avian raptor populations. New infectious diseases associated with high mortality, such as HPAI, might challenge these recent achievements. Avian raptors are especially exposed to pathogens and opportunistic microbiota, when they prey on infected, weakened animals and some species are even scavenging on carcasses. However, hunters and facultative scavengers should have evolved increased resistance to infectious threats from their prey (Zepeda Mendoza et al., 2018; Zou et al., 2021), and even provide beneficial functions by removing potentially infectious carcasses (and their pathogens) from the ecosystem (Plaza, Blanco, & Lambertucci, 2020). Nevertheless, from an evolutionary perspective, HPAIV H5 (gs/GD) is a very recent pathogen in wild birds and yet, no such resistance mechanisms could have been positively selected in avian raptors, including specialized scavengers like vultures (Ducatez et al., 2007). Conceivably, HPAIV H5 infection in immunologically naïve avian raptor species has been shown to induce severe and often fatal disease. These findings have prompted investigations of using avian raptors as indicator species to monitor geographical expansion of HPAIV activity in general and incursion events of HPAIV H5 into new regions, as recently described for the transatlantic spread of HPAIV H5, clade 2.3.4.4b, via Iceland (Günther et al., 2022; Lee et al., 2019).

Our retrospective analysis clearly confirmed a high infection risk for raptors at the end of the food chain, in particular for large to medium-sized avian raptor species, during the epizootic years 2016/17 and since 2020. Thus, the informative status of avian raptors with respect to virological investigations regarding HPAIV is obvious. This became apparent also when analysing the HPAIV H5 genotypes of raptor-born viruses: The HPAI epizootic 2020-2021 in Germany was caused by numerous different subtypes and genotypes of HPAIV H5 (King et al., 2022). Almost a quarter of all different genotypes was also found in various raptor hosts. Their frequency in raptors is proportional to their occurrence in other avian hosts. An exception is genotype Ger-11-21-N1.4 (A/buzzard/Germany-SH/AI07099/2021-like) which has been found exclusively in a single unspecified buzzard. This suggests the origin of Ger-11-21-N1.4 in another primary, avian host, that remained unspecified and undetected, e.g. due to a very localized and restricted occurrence of this virus strain. WTSEs seemed to be particularly informative targets within (passive) surveillance approaches, showing that over time approximately every eighth individual WTSE tested was confirmed positive for HPAIV H5 (Table S1). Indeed, we have been able to provide data to confirm the role of raptor species as suitable indicators for a general HPAI-surveillance and to highlight their importance to reflect even temporal and geographical patterns of genetic variants. During the epizootic 2016/17, juvenile and immature WTSEs have been affected more frequently than (sub)adult ones; nestlings were not affected at all since the virus was only recorded outside the hatching season (Krone et al., 2018). HPAIV H5N1 in a WTSE hatch (n=2) has first been found in the Northern German federal state SH in May 2021. To our current knowledge this is the first detection of HPAIV H5 in nestlings of WTSEs in Germany and matched with a report from Estonia over the same time period (Estonian University of Life Sciences, 19.05.2021). Virological testing during the second year of prospective sampling in MWP, Germany, yielded similar observations of sporadically infected WTSE nestlings (Figure 3) in 2022, when eight nestlings of five locations have been confirmed positive for HPAIV H5, clade 2.3.4.4b.

At the time of sampling, nestlings #77A and #84A did not show severe neurological signs, as described for HPAIV-infected birds before, only a markedly, but unspecific, depression (#84A). In both occasions, another HPAIV-positive nestling was found dead in or in close proximity to the nest. HPAIV RNA was detected in all organ samples taken from these deceased nestlings and confirmed systemic infections in accordance to prior findings during the 2020-21 epizootic by Caliendo, Leijten, et al. (2022).

In contrast to the virological HPAIV testing, few studies have focused on serum antibody analysis. Previous studies failed to detect AIV-reactive antibodies in raptor nestlings during similar sampling approaches in Northern Europe (Gunnarsson et al., 2010; Lee et al., 2019). Here, we analysed raptor sera on a larger scale and detected antibodies against IVA NP (n=8; in 2017, 2021 and 2022) and H5 (n=1; in 2022) exclusively in WTSE nestlings. The fact that we were able to also confirm one H5-seropositive case (#103A; Figure 5-A,) in two independent assays (refer to methods), suggests the reliability of the commercial kits utilized but not validated for raptors due to the lack of reference sera.

NP-antibody detection rates in WTSE nestlings of 5.0% (2021) and 7.9% (2022) appear low given the massive HPAIV H5 outbreak scenarios and the presumed high likelihood of parental female WTSE for exposure during the last two years. Still the single WTSE nestling #103A sampled in 2022 remained the only evidence in a nestling of antibodies against H5 of clade 2.3.4.4 (Figure 5A, Table S5). Due to the highly variable age at which the animals are ringed (and thus sampled), we cannot rule out the possibility that samples were taken, at least in some cases, at a time when maternal antibodies had already declined below detection levels and an active specific immune response had not been generated by the nestling.

Thus, the data may present a vast underestimation of the true seroprevalence in adult female WTSEs. No literature on the stability of maternal antibody levels in WTSEs after hatching exists, but studies among other avian species suggest a rapid decline of maternal antibody levels (van Dijk, Mateman, & Klaassen, 2014; Velarde, Calvin, Ojkic, Barker, & Nagy, 2010), depending on the initial level of yolk-derived antibodies and the respective test sensitivity. Testing younger nestlings closer to hatch might have provided more conclusive results but there is a minimum age of more than five weeks that needs to be considered when sampling is to be combined with ringing.

The case of the WTSE nestling #103A exemplifies the problems of interpretation in a seropostive case: It has been sampled between six to seven weeks after hatching and tested positive for H5-antibodies. Assuming the time period of decreasing maternal antibodies as a matter of a few weeks, not days or months after hatching, the sero-response of nestling #103A cannot be clearly associated with either maternal antibodies or seroconversion after direct contact with HPAIV H5. The latter, unconfirmable, assumption would raise hope that even WTSE nestlings, under certain circumstances, may overcome HPAIV H5 infection.

Sampling of adult birds would have allowed direct measurements of seroprevalence rates; however, adult raptors are accessible for blood sampling on exceptional occasions only. Sampling of large-sized adult raptors is mainly limited to wild bird rescue centers, when birds are admitted for care and such sample set would be skewed by the dominance of samples from non-healthy birds. One out of eight blood sampled non-nestling raptors, other than WTSE, revealed NP- and H5-reactive antibodies (Table S4). The seroconversion of this adult red kite is interpreted as an indication that some raptor species can overcome an AIV H5 infection. Also, a single adult WTSE out of eleven blood sampled adults (9.1%) revealed H5-antibodies, simultaneously the only adult WTSE tested in 2022.

Overall, sampling of nestlings is an elegant option to gain insights into wild avian raptor populations. Although only few trained experts are concerned, direct contact with, as shown here, nestlings shedding HPAIV H5 asymptomatically cannot be excluded. Zoonotic transmission routes via injuries caused by the bird’s beak or claws contaminated with feces or carrion remains can be envisaged when ringing or sampling avian raptors. Strict hygiene measures are required and even cessation of bird ringing activities in confirmed HPAI hotspot regions should be considered to securely prevent spill-over events to humans, but also to avoid bird ringers (unknowingly) becoming vectors between (breeding) locations via contaminated equipment, clothes and shoes. Avian rehabilitation centers and clinics face a similarly high risk of being confronted with HPAIV-H5-positive wild birds. Caliendo, Mensink, et al. (2022) pointed out the importance of increased awareness for those institutions, including continuous education of employees, for adequate quarantine measures to prevent inadvertent spread. Routine virological screening of admitted raptors is highly recommended as exemplified here by the case of WTSE #72A shown here initially misdiagnosed for lead intoxication, one of the most common causes of death in (sub)adult WTSE, but instead being HPAIV H5-infected.

Promoting and maintaining a stable reproduction ratio over many years supported by a high number of breeding pairs is essential for a stable population of long-lived k-strategists such as WTSEs. Reducing this rate by removing adult breeding-competent individuals or by impacting hatching and upbringing of chicks may ultimately lead to population instability. The enzootic status of HPAIV H5 in Germany combines these risks for WTSE as shown here for juvenile to (sub)adult WTSE deaths and five affected hatches in MWP in 2022. Contrary, however, to the expectations and previous reports from bald eagle populations in North America (Nemeth et al., 2023), the overall breeding success rate for this region remained unaffected (Figure 6), even if further cases might have remained undetected. It remains to be determined whether cross-immunity of the parental female birds and, therefore, maternal antibodies in their nestlings might have contributed. Our serological data gave some evidence in that direction.

Furthermore, there is a staggered start of breeding of WTSE pairs within the same region, preventing that all clutches hatch within a very narrow period of time. This would reduce the influence of time-sensitive risk factors to the overall population, mainly related to weather conditions but probably extrapolatable also to the prevalence of pathogens in prey. However, such effect may vary in (coastal) regions where WTSEs have stronger ties to water areas with seabird colonies. The latter have been hit severely by HPAIV H5, culminating even in mass mortalities (Anne Pohlmann et al., 2023). Removal of possibly HPAIV-infected carcasses by human activities has been shown to have positive effects on such colonies (Knief et al., 2023). Yet, this kind of carcass removal can never be as efficacious (and timely) as the scanning activities of birds of prey. In addition, the hunting behaviour e.g. of peregrine falcons cannot be manipulated to be distracted from infected and weakened live prey. Thus, it cannot be excluded that detrimental impacts will develop nevertheless over time in case the enzootic HPAI H5-status in the regional wild bird populations is to continue. With the current continuation of HPAIV H5 circulation in Europe black-headed gulls (*Chroicocephalus ridibundus*) became the dominantly affected species; while no increase in WTSE cases are reported, peregrine falcons and Eurasian eagle owls are found HPAIV H5-infected at increasing rates. In this context, particular adaptations of new emerging genetic variants of gs/GD HPAIV such as the gull-adapted reassortant genotype BB to certain prey species may lead to shifting risks for different raptor species (EFSA (European Food Safety Authority), ECDC (European Centre for Disease Prevention and Control), EURL (European Reference Laboratory for Avian Influenza), Adlhoch, Fusaro, Gonzales, Kuiken, Mirinaviciute, et al., 2023).

## Conclusions

Overall, our results on HPAIV H5 found in raptor species, particularly WTSE, common buzzards, Northern goshawks, peregrine falcons and Eurasian eagle owls, during passive surveillance confirm their suitability as important indicators for the occurrence of the pathogen, including detection of temporally and geographically restricted variants. Prospective screening of avian raptor nestlings revealed the presence of maternal antibodies or seroconversion in WTSE chicks. The still low AIV-seropositivity rate of nestlings in the examined WTSE population indicates a particular risk for naïve nestlings to alimentary HPAIV infections on the nest and after fledging. This became evident by multiple findings of systemic fatal infections in WTSE nestlings in MWP in 2021 and 2022. As yet, no direct detrimental influence on breeding success rates of WTSE was evident in the region. The combination of scientific bird ringing and sampling for disease surveillance seems highly appropriate in terms of coordinated species protection but requires heightened awareness and strict hygiene measures to avoid inadvertent pathogen carryover and human exposure.

## Acknowledgements

We thank Aline Maksimov, Diana Parlow, Mareen Grawe, Cornelia Illing, Sabine Schiller and Kristin Bishop for their excellent technical assistance, as well as Annika Graaf-Rau and Angele Breithaupt for their great support regarding logistics and pathology sessions. We highly appreciate the immense effort of all volunteers participating within bird monitoring and bird ringing activities, including ornithologists, tree climbers and further helpers involved – especially to the bird ringers, Rene Feige, Samuel Knoblauch, Torsten Lauth, Torsten Marczak and Mario Müller. Furthermore, we would like to thank Frank Tetzlaff, who enabled our sampling activities in the wild bird rescue center of the “Tierpark Greifswald” and also supported this work as bird ringer.

## Supportive information

Additional supporting information may be found in the online version of the article at the publisher’s website.

